# Fecal Virome of Southeastern Maned Sloth (*Bradypus crinitus*)

**DOI:** 10.1101/2023.06.23.546151

**Authors:** Amanda Coimbra, Mirela D’arc, Filipe Romero Rebello Moreira, Matheus Augusto Calvano Cosentino, Francine Bittencourt Schiffler, Thamiris dos Santos Miranda, Ricardo Mouta, Déa Luiza Girardi, Isabelle Gomes de Matos, Victor Wanderkoke, Gabriel Medeiros, Monique Lima, Thiago Henrique de Oliveira, Talitha Mayumi Francisco, Flávio Landim Soffiati, Suelen Sanches Ferreira, Carlos Ramon Ruiz-Miranda, Marcelo Alves Soares, André Felipe Andrade dos Santos

## Abstract

We report a viral metagenomic analysis of fecal samples from *Bradypus crinitus*, a recently described sloth species that occurs in the Atlantic Forest of Espírito Santo and Rio de Janeiro states, Southeast Brazil. Through Illumina sequencing, we generated a total of 2,065,344 raw reads, of which 945,386 reads (45.77%) passed the quality and size filter. The highest proportion of them was assigned to Eukarya, followed by Bacteria and only a small proportion to Virus. However, we identified 24 viral families using distinct taxonomic assignment tools, including phages and vertebrate viruses, such as retroviruses and papillomaviruses. Also, we identified four bacterial genus already associated with disease in sloths. Our study sheds light on the microbiome of a previously unexplored species, further contributing to the comprehension of metagenomic global diversity.

## MAIN TEXT

The maned sloth is endemic to the Brazilian Atlantic Rain forest and has been categorized as vulnerable by the International Union for Conservation of Nature (Chiarello and Moraes-Barros, 2013). A recent taxonomic revision divided the group into two species: northeastern maned sloth (*Bradypus torquatus*), occurring in the states of Bahia and Sergipe; and southeastern maned sloth (*Bradypus crinitus*), in the states of Espírito Santo and Rio de Janeiro, respectively (Miranda *et al*., 2023). Serology studies elucidated that northeastern collared sloths are well known reservoirs of several arboviruses genera: *Orthobunyavirus*, as Utinga virus (UTIV) (Seymour, 1985) and Caraparu virus (CARV) (Catenacci *et al*., 2018); *Flavivirus*, as Dengue virus (DENV 1 to 4) (Catenacci et al., 2018), Rocio virus (ROCV) (Catenacci et al., 2018), Bussuquara virus (BSQV) (Catenacci *et al*., 2018), Saint Louis encephalitis virus (SLEV) (Seymour, 1985), Ilheus virus (ILHV) (Seymour, 1985) and Yellow fever virus (YFV) (Catenacci *et al*., 2018); and also, *Alphavirus*, as Eastern equine encephalitis virus (EEEV) (Catenacci *et al*., 2018). However, apart from arbovirus, little is known about the natural viral diversity of maned sloths. Recent advances in high-throughput sequencing (HTS) technologies have allowed comprehensive access to the virosphere, helping to fill gaps in the diversity of eukaryotic viruses (Zhang *et al*., 2018; Harvey and Holmes, 2022). There are only a few studies that used this technique to assess the enteric virome of sloths (Dill-McFarland *et al*., 2016; Rojas-Gätjens *et al*., 2022). Here, we describe the viral diversity obtained by HTS from fecal samples from southeastern maned sloths, collected in the Atlantic Forest of Rio de Janeiro.

The study was carried out between November 2018 and July 2019 in the localities of *Igarapé* and *Dois irmãos* Farms, in the state of Rio de Janeiro (**Table 1**). Through active search, adult and subadult healthy animals were located and captured directly from the treetops. Fecal samples were collected from seven southeastern maned sloths at the moment of animal’s manipulation and transferred to 50 mL falcon tubes with a volumetric ratio roughly 1:1 of RNA*later*(tm) (Thermo Fisher Scientific, Walham, USA). Samples were kept at room temperature and shipped to the *Laboratório de Diversidade e Doenças Virais* (LDDV) at *Universidade Federal do Rio de Janeiro* (UFRJ), Rio de Janeiro, Brazil, to be stored at -80 °C. For the virome protocol, the samples were vigorously vortexed until complete homogenization. Then, approximately 1 mL of sample was transferred to the extraction bead tube (MP Biomedicals, CA, USA) to break up the debris. The supernatant was collected from each sample and an equimolar volume was pooled in one single tube to maximize sequencing efficiency. The next step was Illumina library construction, following the general CDC protocol methodology (Kohl *et al*., 2015) with previously described modifications (Cosentino *et al*., 2022). HTS was conducted on an Illumina MiSeq platform using the MiSeq V2 cartridge with pair-ended 2×151 cycles. Sequencing data was processed with a custom pipeline, also previously described (Cosentino *et al*., 2022). To avoid false positive results due to sequencing index-hoping (incorrect assignment of reads to a given sample) (Farouni *et al*., 2020), we considered as invalid the identification of a viral family when it was detected by a number of reads that was less than 1% of the highest count identified for the same family among all other libraries sequenced in the same run.

**Table 1:** General information of collected specimens.

| **ID** | **Date** | **Location** | **GPS\*** | **Age** | **Sex** |
| --- | --- | --- | --- | --- | --- |
| BT01 | Nov/2018 | Igarapé, Silva Jardim, RJ | -22.50713, -42.30954 | Subadult | F |
| BT03 | July/2019 | Dois irmãos, Silva Jardim, RJ | -22.38538, -42.27638 | Subadult | M |
| BT05 | June/2019 | Igarapé, Silva Jardim, RJ | -22.42530, -42.02737 | Subadult | M |
| BT06 | June/2019 | Dois irmãos, Silva Jardim, RJ | -22.50713, -42.30954 | Adult | F |
| BT07 | July/2019 | Dois irmãos, Silva Jardim, RJ | -22.38538, -42.27638 | Adult | F |
| BT08 | July/2019 | Dois irmãos, Silva Jardim, RJ | -22.38538, -42.27638 | Adult | F |
| BT09 | July/2019 | Igarapé, Silva Jardim, RJ | -22.38538, -42.27638 | Subadult | F |

\*Global Positioning System in Decimal Degrees format.

The pooled samples generated a total of 2,065,344 raw reads, of which 945,386 reads (45.77%) passed the quality and size filter. Taxonomic assignments were performed with both Kraken2 (Wood and Salzberg, 2014) (standard database) and Diamond v.2.0.14 (Buchfink *et al*., 2015) (NCBI nr database). Using Kraken2, a total of 368,564 reads (38.98%) were classified, being the highest proportion assigned to Eukarya (253,144 reads; 68.7%), followed by Bacteria (113,958 reads; 30.9%) and only a small proportion to Virus (318 reads; 0.09%) (**Supplementary Table 1**). Among the viral reads, 24 families were identified (**Figure 1**; **Supplementary Table 2**). Diamond classified 98,694 reads (10.44%), the majority were bacterial (54,910 reads; 55.6%), followed by eukaryotes (42,975 reads; 43.5%) and, similar to Kraken2, a small proportion of virus (294 reads; 0.3%) (**Supplementary Table 1**). By analyzing the sampled bacteriome, we were able to identify bacterial reads already identified in sloths such as *Bordetella, Citrobacter, Escherichia, Salmonella, Coxiella, Anaplasma* and *Ehrlichia* (**Supplementary File 1**),of which the first three and some strains of the fourth were able to cause disease in these animals (Smith and Ruple, 2021).

**Figure 1:**
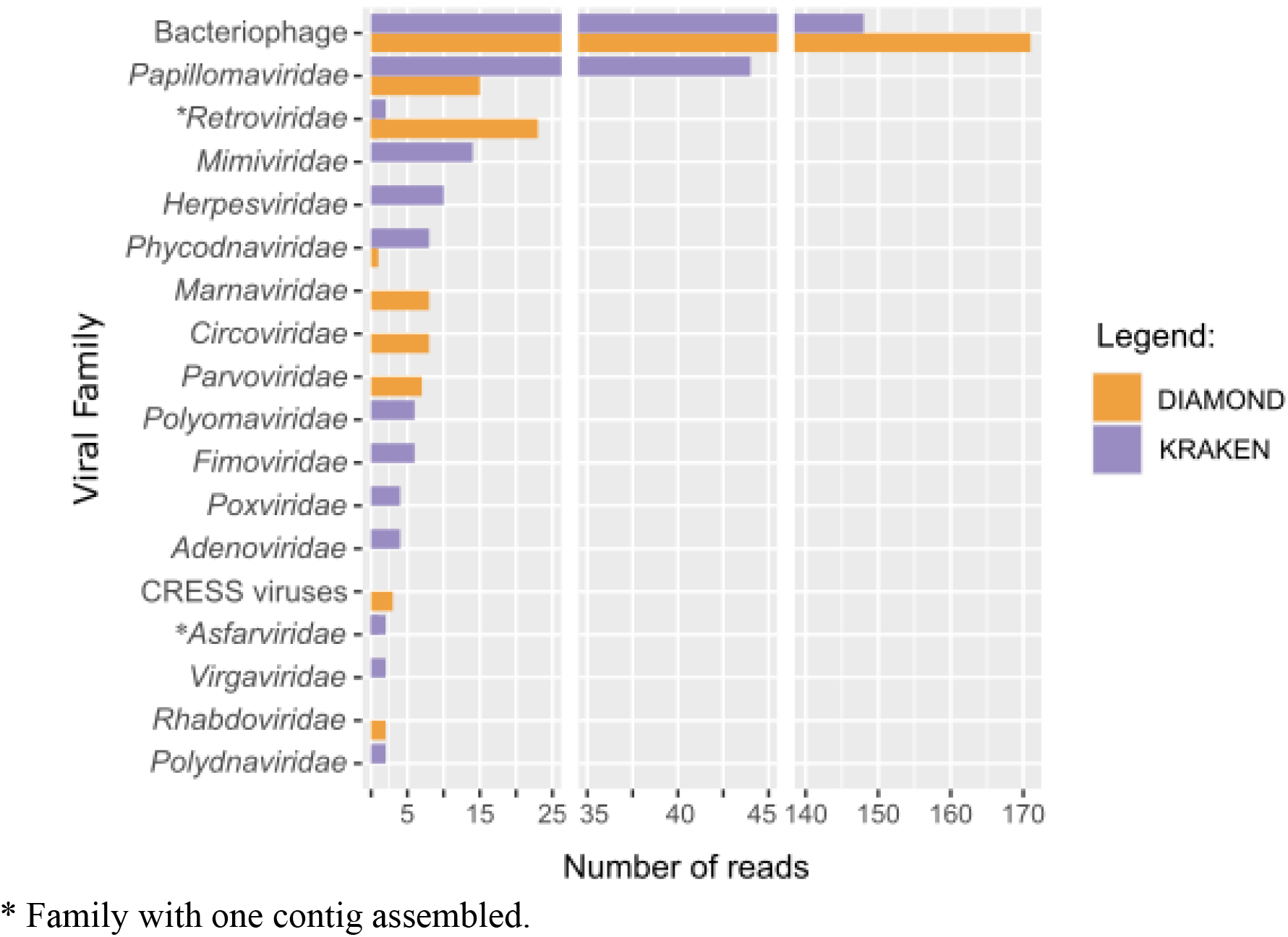
Viral families identified in sloth fecal samples.

A total of 15 viral families were identified by Diamond, and nine (60%) were in agreement with Kraken2. Among the viral reads validated by both tools, the bacteriophages were the most abundant (*Autographiviridae, Microviridae, Myoviridae, Podoviridae, Salasmaviridae, Schitoviridae* and *Siphoviridae*), followed by vertebrate viruses, including sequences from *Papillomaviridae* and *Retroviridae*, commonly linked with animals and humans diseases (**Figure 1**; **Supplementary Table 2**). We also performed *de novo* assembly with MetaSpades v.3.15.3 (Nurk *et al*., 2017) and obtained a total of 5,787 contigs (ranging from 125 - 6,752 nt, median size: 435 nt), with only two viral families identified (*Asfarviridae* with one contig classified by Kraken2 and *Retroviridae* with one contig classified by Diamond). Other viruses related to plants, phytoplankton and protozoa were also assigned, as *Virgaviridae, Marnaviridae* and *Mimiviridae*, respectively. Finally, a large amount of reads (Kraken2: n=576,822; Diamond: n=846,692) were not assigned, which potentially includes new and distantly related viral groups yet to be classified (**Supplementary Table 1)**. The available literature on sloths viral diseases mostly assesses their serological status, thus focusing on past exposure (Catenacci *et al*., 2018; Seymour, 1985). Our study shows the occurrence of current viral infections in sloth populations, also revealing the composition of their largely unknown fecal microbiome, as well as the presence of potentially pathogenic bacteria, and further contributes to the understanding of microorganism diversity. However, we emphasize the need for further studies on this topic, especially regarding its symbiont bacterial composition, which may be associated with the ability of this species to digest its food sources.

## DATA AVAILABILITY

Raw sequencing reads are available under BioProject accession number PRJNA971998 and NCBI SRA accession number SRR24525879. The Supplementary File is available as a separated pdf. Assembled contigs, taxonomic assignment files and Krona plots are available on the project GitHub page (link: https://github.com/lddv-ufrj/sloth_virome).

## ACKNOWLEDGMENTS

Field work was supported by funds from PETROBRAS (process 2017/00606-7) to Carlos R Ruiz-Miranda for the project *Avaliação do efeito de faixas de dutos na conectividade da paisagem para a Mastofauna e análise da eficácia de estruturas de travessias de fauna*. This work also was supported by the Conselho Nacional de Desenvolvimento Científico e Tecnológico/CNPq (grant number 313005/2020-6 to AFS) and Fundação de Amparo à Pesquisa do Estado do Rio de Janeiro/FAPERJ (grants numbers E26/201.193/2022, E26/211.355/2021, and E26/211.040/2019 to AFS). We acknowledge the use of Biology Molecular equipments from *Laboratório de Uso Comum e Apoio Técnico* from *Universidade Federal do Rio de Janeiro* (Rio de Janeiro, Brazil) and all the workers for assistance with sample collection, in special to Caique Ferreira Amaral Soares.

We declare no conflicts of interest. The animals were captured and samples manipulated under the approval and legal consent of the Brazilian Federal Authority (numbers 67274-8 and 64635-5).

