## Supplementary File for "Fecal Virome of Southeastern Maned Sloth (*Bradypus crinitus*)"

**Supplementary Table 1:** Overview data of filtered reads (n=945,386 corresponding to 45.77% of the total sequenced) and assembled contigs (n=5,787) taxonomically classified by Diamond and Kraken2.

| | | Diamond | Kraken2 |
|---|---|---|---|
| No hits | Reads (%) | 846,692 (89.7%) | 576,822 (61%) |
| | Contigs (%) (min - max) | 3,392 (58.6%) (125 -1,840 nt) | 4,577 (79%) (125 - 4,162 nt) |
| Mapped reads | | 98,694 (10.4%) | 368,564 (38.9%) |
| Mapped contigs (min - max) | | 2,395 (186 - 6,752 nt) | 1,210 (129 - 6,752 nt) |
| Bacteria | Reads (%) | 54,910 (55.6%) | 113,958 (30.9%) |
| | Contigs (%) (min - max) | 508 (21.2%) (375 - 3,780 nt) | 511 (42.2%) (131 - 4,802 nt) |
| Eukaryota | Reads (%) | 42,975 (43.5%) | 253,144 (68.7%) |
| | Contigs (%) (min - max) | 1,883 (78.6%) (186 - 6,752 nt) | 683 (56.5%) (129 - 6,752 nt) |
| Viral | Reads (%) | 294 (0.3%) | 318 (0.09%) |
| | Contigs (%) (min - max) | 2 (0.08%) (381 - 1,441 nt) | 1 (0.08%) (409 nt) |
| Others/ Unassigned | Reads (%) | 515 (0.52%) | 1,144 (0.31%) |
| | Contigs (%) (min - max) | 2 (0.08%) (437 - 445 nt) | 15 (1.24%) (186 - 735 nt) |

**Supplementary Table 2:** Viral families classified by Kraken2 and Diamond.

| VIRUS FAMILY | KNOWN HOSTS | READS K / D* | CONTIGS K / D* |
|---|---|---|---|
| ***Adenoviridae***** | **Vertebrates** | 4 / 0 | 0 / 0 |
| ***Asfarviridae*** | **Vertebrates** \| Invertebrates | **2 / 0** | **1 / 0** |
| *Autographiviridae* | Bacteria | 6 / 6 | 0 / 0 |
| ***Circoviridae*** | **Vertebrates** | **0 / 8** | **0 / 0** |
| **CRESS viruses** | **Vertebrates** \| Environmental | **0 / 3** | **0 / 0** |
| ***Fimoviridae*** | **Vertebrates** | **6 / 0** | **0 / 0** |
| ***Herpesviridae*** | **Vertebrates** | **10 / 0** | **0 / 0** |
| *Marnaviridae* | Phytoplankton | 0 / 8 | 0 / 0 |
| *Microviridae* | Bacteria | 0 / 14 | 0 / 0 |
| *Mimiviridae* | Protozoa | 14 / 0 | 0 / 0 |
| *Myoviridae* | Bacteria | 34 / 46 | 0 / 0 |
| ***Papillomaviridae*** | **Vertebrates** | **44 / 15** | **0 / 0** |
| ***Parvoviridae*** | **Vertebrates** \| Invertebrates | **0 / 7** | **0 / 0** |
| *Phycodnaviridae* | Algae | 8 / 1 | 0 / 0 |
| *Podoviridae* | Bacteria | 6 / 22 | 0 / 0 |
| *Polydnaviridae* | Invertebrates | 2 / 0 | 0 / 0 |
| ***Polyomaviridae*** | **Vertebrates** | **6 / 0** | **0 / 0** |
| ***Poxviridae*** | **Vertebrates** | **4 / 0** | **0 / 0** |
| ***Retroviridae*** | **Vertebrates** | **2 / 23** | **0 / 1** |
| ***Rhabdoviridae*** | **Vertebrates** \| Invertebrates \| Plants | **0 / 2** | **0 / 0** |
| *Salasmaviridae* | Bacteria | 2 / 3 | 0 / 0 |
| *Schitoviridae* | Bacteria | 4 / 8 | 0 / 0 |
| *Siphoviridae* | Bacteria | 96 / 72 | 0 / 0 |
| *Virgaviridae* | Plants | 2 / 0 | 0 / 0 |

*K = Kraken2 and D = Diamond.

** Featured data on viral families that infect vertebrates.
